# Pursuing the limits of child survival in the most and least developed countries

**DOI:** 10.1101/591925

**Authors:** Iván Mejía-Guevara, Wenyun Zuo, Laust H. Mortensen, Shripad Tuljapurkar

## Abstract

The epidemiological transition from young to old deaths in high-income countries reduced mortality at all ages, but a major role was played by a decline of infant and child mortality from infectious diseases^1,2^ that greatly increased life expectancy at birth^2,3^. Over time, declines in infectious disease continue but chronic and degenerative causes persist^4,5^, so we might expect under-5 deaths to be concentrated in the first month of life. However, little is known about the age-pattern of this transition in early mortality or its potential limits. Here we first describe the limit using detailed data on Denmark, Japan, France, and the USA— developed countries with low under-5 mortality. The limiting pattern of under-5 deaths concentrates in the first month, but is surprisingly dispersed over later ages: we call this the early rectangularization of mortality. Then we examine the progress towards this limit of 31 developing countries from sub-Saharan Africa (SSA)—the region with the highest under-5 mortality^6^. In these countries, we find that early deaths have large age-heterogeneities; and that the age patterns of death is an important marker of progress in the mortality transition at early ages. But a negative association between national income and under-5 mortality levels, confirmed here, does not help explain reductions in child mortality during the transition.

## Main

The epidemiological transition from young to old deaths in high-income countries has characterized mortality decline at all ages. But the decline of infant and child mortality from exogenous causes (infections or parasites, accidents)^1,2^ led to substantial increases in life expectancy at birth^2,3^, and an increasing modal age-at-death for adults^7^. Over time, as declines in exogenous mortality continue, under-5 mortality (U5M) should shift towards ages close to birth, but is likely to remain high during the first month of life due to the persistence of endogenous causes of death (congenital malformations, injuries connected with birth)^8^. Here we identify a limit reached by the rich countries with very low U5M, and call this limiting pattern of death the *Early Rectangularization of the Mortality Curve*. This limiting age-distribution for U5M in rich countries is not reflected in Sustainable Development Goal 3, which sets a target of neonatal (NMR) and under-5 mortality rates of 12, respectively 25, deaths per 1,000 births by 2030 for countries with currently high U5M^6^. Here, we investigate the extent of progress towards our U5M limit in 31 sub-Saharan African (SSA) countries.

In any year, U5M levels vary dramatically with age, and change at different rates with respect to age and across countries. These factors are not accounted for by established techniques, e.g., Gini coefficient, standard deviation of age at death, and life disparity (these do work at old ages as measures of mortality compression or rectangularization^1,9,10^, or as measures of lifespan inequality at adult ages^11–14^). To overcome these limitations for U5M, we use a generalized Gini (G_[0]_) index that is proportional to the average mortality rates, and accounts for relative as well as absolute differences across ages (Methods) <u>(1)</u>. We expect inequality as measured by G_[0]_ to decline over time in response to declines in U5M at any age. In contrast, at adult ages inequality in lifespan can decline or increase depending on the ages at which mortality declines^15,16^.

We rely on historical data from Denmark (1901-2016), France (1975-2016), Japan (1980-2016), and the USA (1968-2016), with high-quality and fine-grained U5M records measured in weeks/months (Supplementary Methods). These countries are not only highly developed economically, but also display the lowest U5M levels ever recorded after more than 100 years of continuing declines (Fig 1a). Long-run convergence is shown by U5M in Denmark during the first 80 years of the 20^th^ Century, with G_[0]_ declining from 1.6 to 0.1 x1000 (an average annual decline of 2.4%). More recent data reveal that Japan has reached the lowest concentration of under-5 mortality in 2016 (G_[0]_ = 0.025 x1000) and we argue that Japan is close to the actual limit of U5M at that time as it is converging more rapidly than other developed countries—G_[0]_ in Japan declined by 74% between 1980 and 2016, followed by France (65%), Denmark (60%), and the U.S. (54%) during the same period. The lower level and more rapid decline of neonatal mortality in Japan during the period explains its leading role in the mortality convergence (Fig 1b). By 1980, Japan’s neonatal and under-5 mortality were around one third of the levels set by the SDG-3 by 2030 (4.1 and 8.4 per 1,000 births, respectively), and continued declining to lower levels of 0.7 and 2.3 deaths per 1,000 births by 2016, respectively. Likewise, by 1975 France’s NMR (7.6) and U5M (13.9) were around two-thirds of the respective SDG-3 limits, Denmark reached those limits between 1955-1965 (U5M was 24.7 x1000 in 1955 and NMR was 12.5 x1000 in 1965), and the USA was about to reach them around 1968 (NMR=13.6, U5M=21.5) (Fig. 1b).

**Fig. 1.**
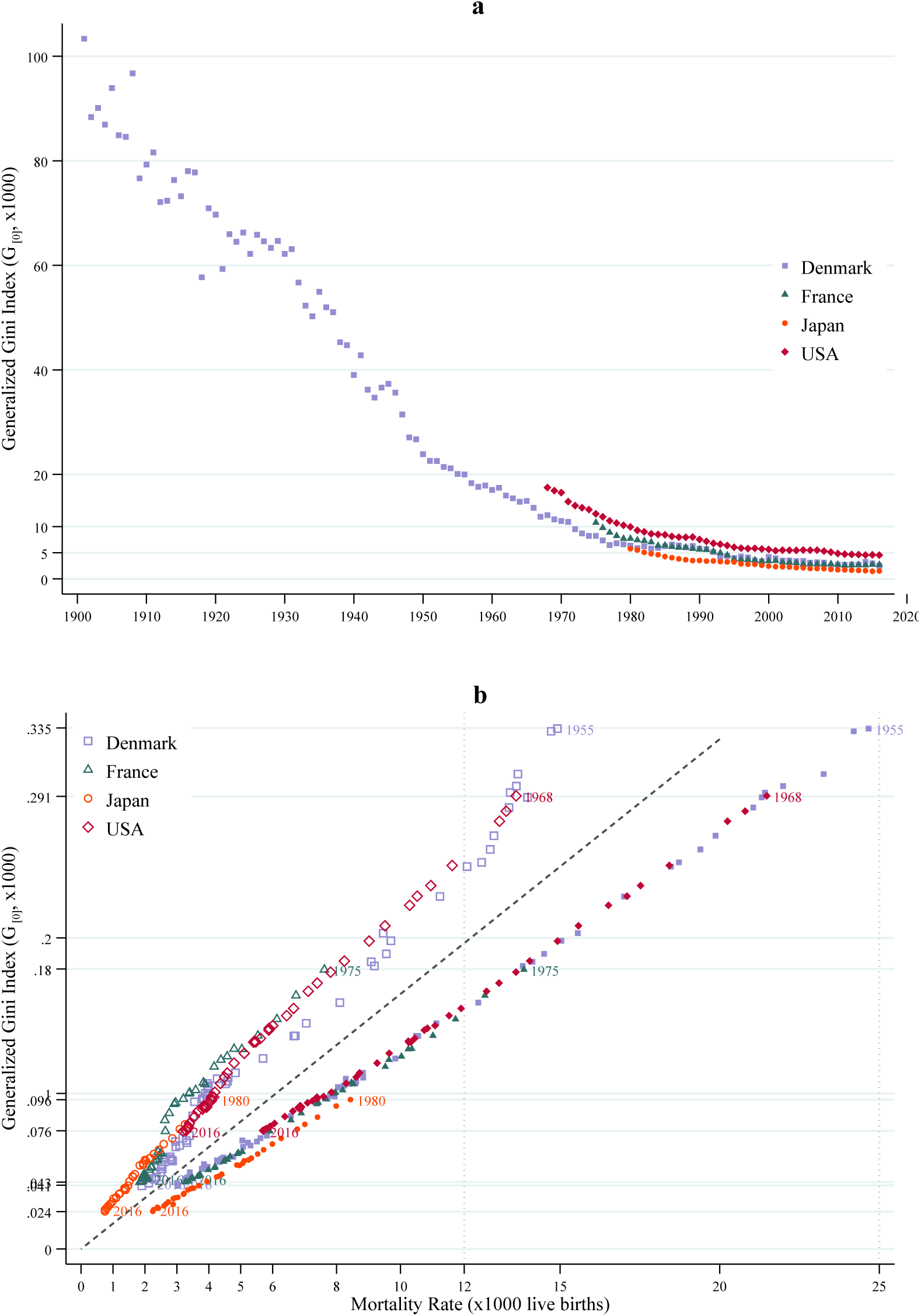
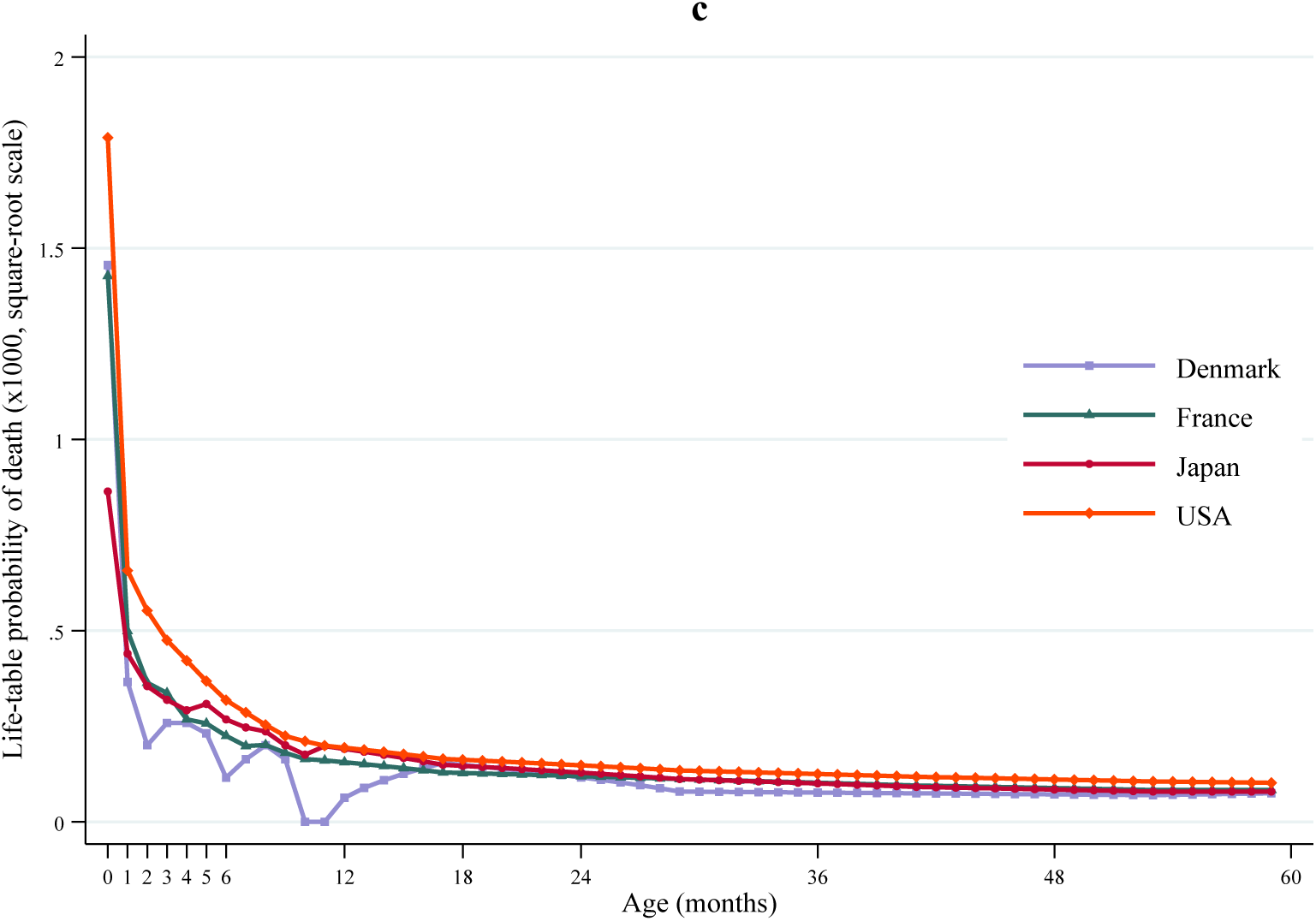
Convergence of under-5 mortality and the limiting patterns of under-5 deaths. **a**, Trends of under-5 mortality (U5M) compression by country over time. **b**, Convergence of Neonatal and U5M in the last 65 years. **c)**, Life-table distribution of under-5 deaths for Denmark and Japan in 2016. Data are for Denmark, France, Japan, and the USA for the period between 1980 and 2016. In **a** and **b**, Mortality compression/convergence was assessed using the generalized Gini index (G_[0]_), which accounts for both changes in the age distribution and levels of mortality across countries over time, and smaller values of G_[0]_ indicate early compression (less concentration of deaths), less inequality, or more rapid convergence of under-5 mortality. The dashed diagonal line in **a** illustrates the hypothetical case of perfect rectangularization of the U5M curve where all deaths are concentrated at age 0 (the mean probability of death 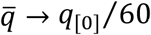, where *q*_[0]_ represents the neonatal mortality rate). In **c**, we illustrate the early rectangularization of the mortality distribution, represented the empirical limiting patterns of under-5 deaths in these four countries in 2016 (probabilities of death are per 1,000 in a squared-root scale).

In Figure 1c we show that the limiting pattern of under-5 mortality can vary, but the rectangular (or ‘L’) shape—characterized by an age pattern where mortality rates decline overall but remain high during the first month of life— remains a distinctive characteristic even at very advance stages of the mortality transition. Had all deaths occured within the first month of life (as illustrated by the dashed diagonal line in Fig. 1b), the limiting pattern would be represented by a vertical line at age 0 and flat for older ages. The recent age-pattern in Japan is much less concentrated, even though U5M remains high in the first month after birth. The recent age-patterns of U5M from Denmark and France are similar to that in Japan, but with higher NMRs; whereas the age pattern from the USA also exhibits an ‘L’ shape, but neonatal and infant mortality (deaths within the first year of life) rates were the highest among the 4 countries (Fig. 1c).

The speed of convergence to the limiting pattern is determined by the combined effects of changing mortality levels and age structure. Thus, with some variations in neonatal mortality levels, Denmark and France have experienced similar transitions in the shape and concentration of mortality during the last 40 years (Fig. 2), but both are still behind Japan. The transition in the USA has been slower as its age pattern in 2016 was still similar to that observed in Japan in 1980, which explains the remaining differences in mortality concentration and age-inequality between these countries during the last 40 years. The generalized Lorenz curve flattens rapidly in Japan as its G_[0]_ approaches the mean under-5 mortality rate (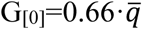; i.e., closest to the theoretical limit of 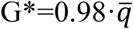), with the lowest 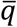 observed among these 4 countries (at 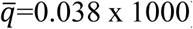) (Fig. 2b).

**Fig. 2.**
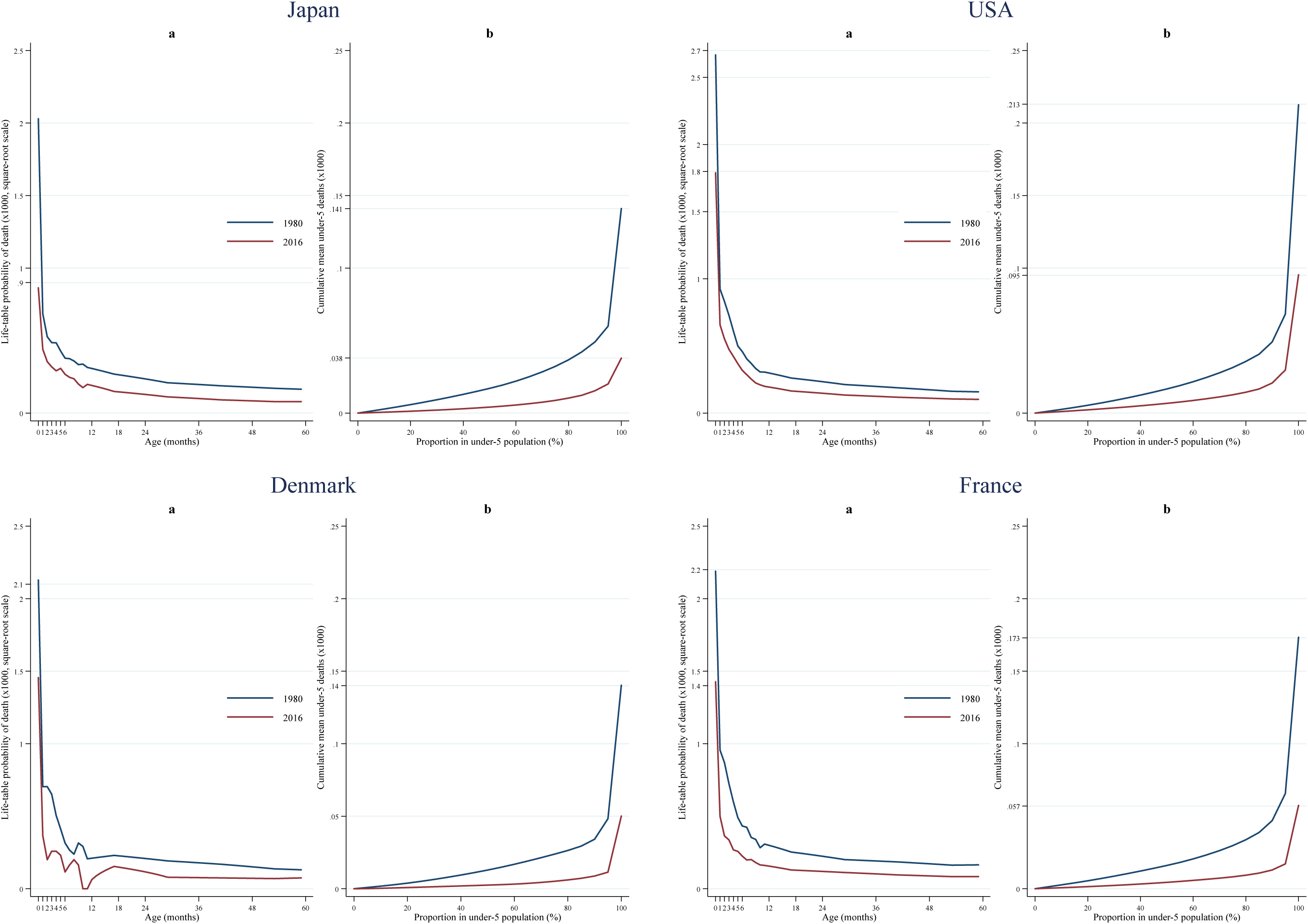
Age patterns of under-5 deaths and inequality in the age at death. **a**, Detailed age-at-death distribution of under-5 deaths in four developed countries (Denmark, France, Japan, and the USA). Age is measured in months in the x-axis and life-table probabilities of dying are represented in the y-axis (x1000 in a square-root scale). **b**, Generalized Lorenz curves in the same selected countries. Generalized Lorenz curves are constructed as the product of the standard Lorenz curve (Extended Data Fig. 1) and the mean mortality rates across ages (q^−^, measured in months). It therefore illustrates the cumulative percentage of under-5 population (x-axis), ranked by the level of mortality at each age, and graphed against the cumulative mean under-5 deaths (y-axis). The Lorenz curve flattens down more rapidly in Japan as its G_[0]_ approaches the mean under-5 mortality rate (G_[0]_=0.66·q^−^; i.e., closer to the theoretical limit of G*=0.98·q^−^), with the lowest q^−^ observed among these 4 countries (at q^−^=0.038 × 1000) (Fig. 2b) (Supplementary Methods and Extended Data Table1).

Heterogeneities in mortality shapes, levels, and rates of convergence are far more pronounced across the least developed countries, as their epidemiologic transition has started more recently. To appreciate this from a historical perspective, we note that Denmark in 1901 exhibited high mortality for neonates and infants and then declining smoothly and rapidly with age. But by 2016, U5M in Denmark exhibits a clear rectangular shape (Extended Fig. 1). We observe roughly similar declines for Rwanda between 1994 and 2014, with a rapid transition and early rectangularization — even so, although Rwanda is likely to reach the SDG-3 by 2030^17^, the country is likely to be well short of the U5M limit in developed countries even by 2050 (Extended Fig. 1). In general, the SSA region is converging unevenly, as shown by values of the generalized Gini (Fig. 3a). As we show there, Rwanda and Senegal have made outstanding progress in mortality decline and reduced age-inequality during the last 20 years; meanwhile, countries like Nigeria have fallen behind and still exhibit mortality levels and shapes only observed in the developed world a century ago.

**Fig. 3.**
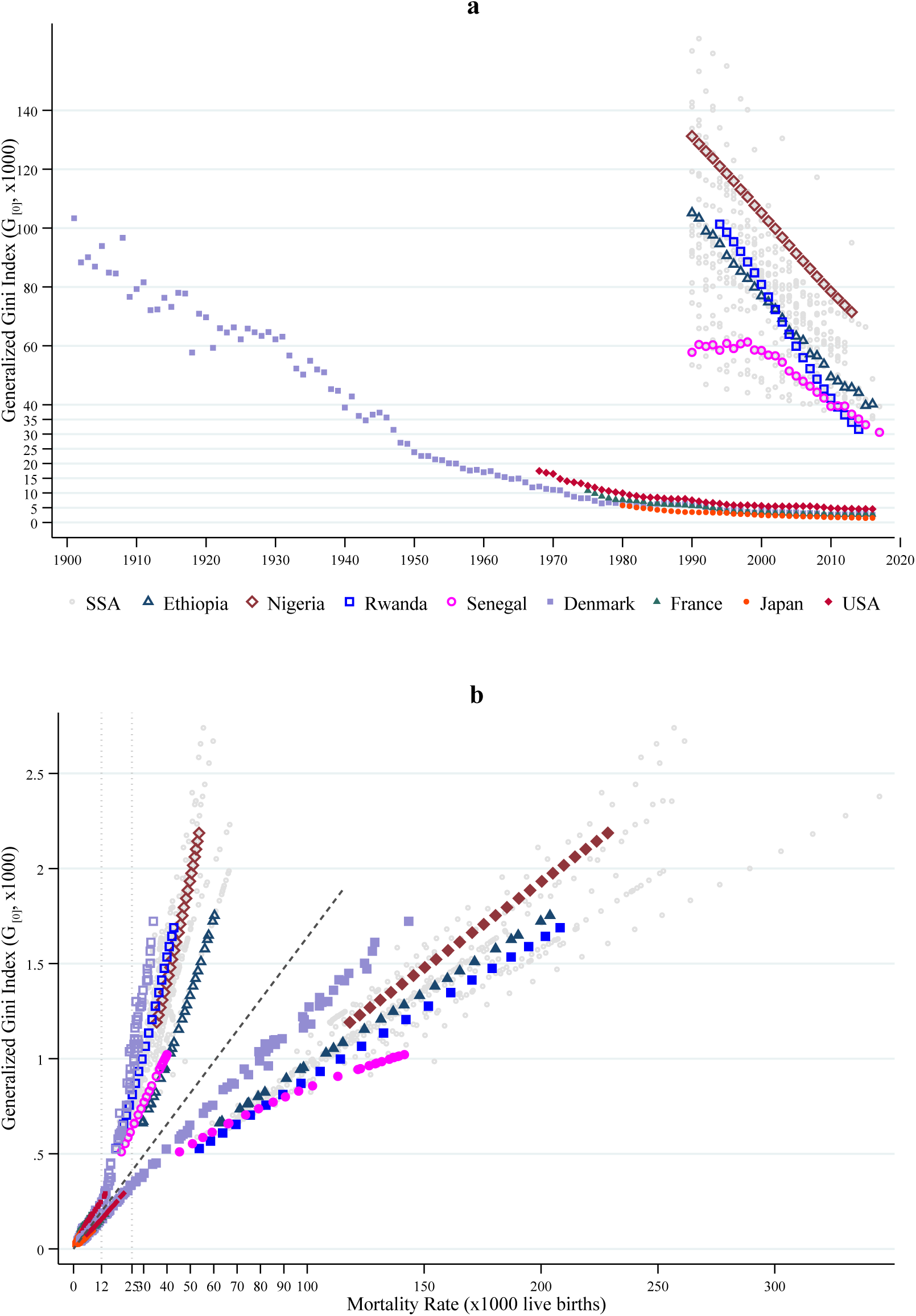
Compression and convergence of under-5 mortality. **a**, Trends of under-5 mortality compression by country over time. **b**, Convergence of neonatal (hollow markers) and under-5 mortality (solid markers)by country over time. Data are for Denmark (1901-016), France (1975-2016), Japan (1980-2016), the USA (1968-2016), and from 31 SSA countries (1990-2017). Compression and convergence of mortality were assessed using the generalized Gini index (G_[0]_), where smaller values of G_[0]_ indicate early compression (concentration of deaths towards age 0), convergence to lower mortality levels, or less inequality. Data are from publicly available mortality records for high-developed countries and from the Demographic and Health Surveys for 31 SSA countries (Supplementary Methods).

Unlike Rwanda, other SSA countries will fall short in meeting the SDG-3 targets by 2030. For instance, Kenya would make it by 2050, but Nigeria (and other countries) is expected to fall short even by 2050 (Extended Data Fig. 1)^17^. A major cause of this delayed transition is the slow convergence of neonatal deaths in most countries of the region (Fig. 3b).

At that current pace, what are the prospects of convergence towards Japan’s current limit distribution for countries in SSA? According to our predictions, the generalized Gini in Rwanda and Kenya would still be 10 times larger in 2030 and 2050 than Japan in 2016, respectively; or 19 times larger in Nigeria by 2050 (Extended Data Fig. 1). Therefore, most SSA countries face an important challenge just to meet the SDGs, but convergence to our early rectangularization limit of U5M does not look realistic in the foreseeable future (Fig. 3b). In any case, we expect that the latter convergence will become increasingly challenging in SSA countries, and will require progress in both technology and socioeconomic conditions.

Our analysis relied on data from countries at opposite sides of the income distribution, and by using 2017 mortality and per capita GDP data we found an expected inverse association^18^—a concave-up shape with a declining monotonic hazard—as levels of U5M fall with economic development (Fig. 4a and Extended Data Fig. 2). However, income does not explain the remarkable differences in the rate of mortality decline within economic regions over time (between 1990-2017), particularly at the lower tail of the income distribution—between 1990 and 2017, the association between income and the decline in the under-5 mortality rate changed between 0.20% and 0.73% in low-income countries (LIC), while it was positive and increasing for upper middle-income countries (UMC) or only varied from 0.13% to 0.42% in high-income countries (HIC) (Fig. 4b). These results contribute to the mixed evidence on the relationship between income and health/mortality, and the shape of that association^19,18,20–23^.

**Fig. 4.**
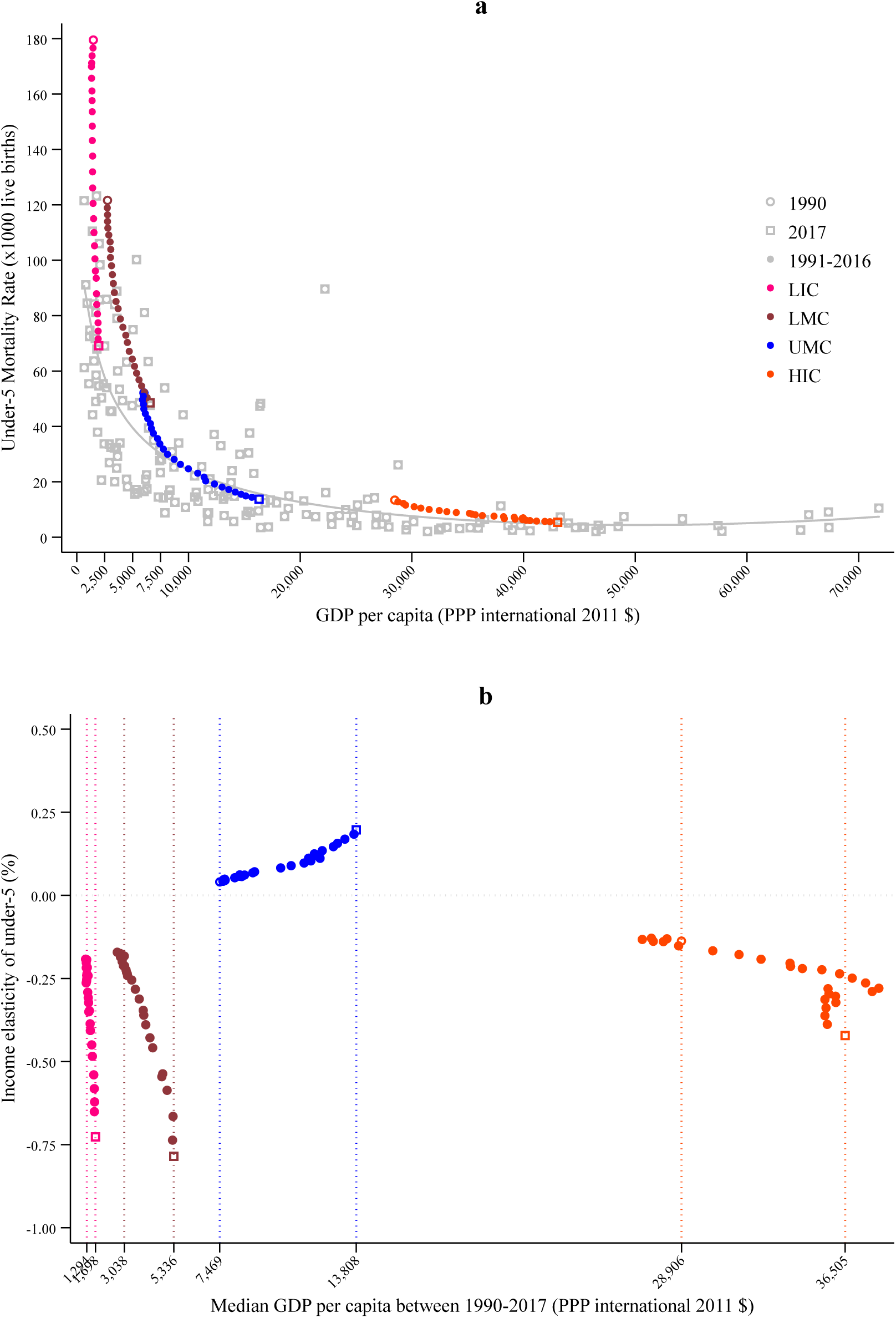
Worldwide association of under-5 mortality and economic development. **a**, Country-wise association of GDP per capita income and under-5 mortality rates in 2017. GDP per capita is expressed in international 2001 Purchasing Power Parity (PPP) United States Dollars (USD), under-5 mortality rates as deaths per 1,000 live births, and yearly average values are shown for the period between 1990 and 2017 by economic region (in color). The adjusted curve represents the prediction of under-5 mortality from an estimated fractional polynomial of income. **b**, Conditional marginal effects of income between 1990 and 2017 by economic region. Income elasticity of mortality represents a % change in under-5 mortality for a % change in GDP per capita. Vertical dotted lines indicate the median GDP at the initial and end of the period in each economic region. Data are from World Development Indicators, retrieved online from The World Bank (2019), which included 217 countries worldwide (8 countries with GDP above 70,000 are not displayed in panel **b**, classified in 4 economic regions: low-income (LIC—pink), low-middle-income (LMC— maroon), upper-middle-income (UMC—blue), and high-income (HIC—orange) (Supplementary Methods).

Our findings also suggest that economic performance was not a precondition for reducing U5M in SSA during its early transition, where policies and interventions have been effective in countries that have met the Millennium Development Goals for child mortality reduction (e.g., Rwanda and Tanzania)^24^. It is less clear, however, whether further progress would be feasible in the region while maintaining low levels of economic development.

Our limit for child survival has scientifically important implications for the causes of early death. It is important in terms of whether future medical/technological advances would allow further reductions of neonatal deaths attributed to endogenous etiology (e.g., conditions originating in the perinatal period or congenital causes)^25^. This limit also has important policy implications for establishing realistic benchmarks in specific contexts; e.g., policies aiming to reach specific goals in poor regions, or to reduce gaps among disadvantaged or minority groups in high-income areas^26^. Our limit also shows that the age distribution of mortality is important in analyzing mortality convergence, and reveals sharp heterogeneities across and within regions. Age-specific progress in SSA may require a combination of structural investments that were effective in high-income countries independently of vaccination and medical technologies^27^, and a significant scale-up of programmes and interventions that target maternal and child care and services^28^. Finally, limited resources in SSA suggest that cost-effective/efficient strategies, such as investments skewed more to primary care than secondary/tertiary care, are likely be more beneficial at the current stage of their mortality transition^29^.

## Methods

### Sources and retrospective mortality data

We collected age profiles of under-1 mortality from publicly available sources from Denmark (1901-2016)^30^, France (1975-2016)^31^, Japan (1980-2016)^32^, and the USA (1968-2016)^33^. Mortality rates for ages 1, 2, 3, and 4 were not available for these 4 countries, but we obtained the corresponding rates as reported in the Human Mortality Database (HMD) for each country^34^ (Supplementary Fig. S1). We then adjusted monthly-based under-5 mortality profiles using a two-dimensional linear interpolation method^35^.

Data from sub-Saharan Africa (SSA) were obtained from birth histories of 106 nationally representative Demographic and Health Surveys (DHS) from 31 countries from the period between 1990 and 2017. The DHS Program collects health and demographic information for women in reproductive age (15-49 years old) and their children, using a two-stage stratified cluster sampling design that defines strata by region and by rural-urban within each region. A sampling frame consisting of census enumeration units or tracks or Primary Sampling Units (PSUs) that cover the entire country is used for the random selection of PSUs in the first stage, with probability proportional to sampling size. Clusters correspond to selected or further split PSUs (or blocks). In the second stage, households are selected systematically from a list of previously enumerated households within each selected cluster or block.

Mortality profiles by age were assessed using full birth history (FBH) data available for individual women aged 15 to 49 years in DHS surveys, where the respondent mother is asked about the date of birth of each of her ever-born children, and the age at death if the child has already died^36^. The complete list of countries included in this study are in the Extended Data Table 1.

### ‘Conditional’ life-table age distribution of under-5 deaths

We used life-table age distributions of death, constructed by Mejía-Guevara et al. (2019) based on the following demographic analysis^17^.

First, using death reports by households and retrospective information from full birth history (FBH) data from DHS^36^, estimates of age-specific death rates m_[x]_ ([x] stands for age in months) were obtained from survey estimates of the number of events and total time to event^36^, and used to compute period life-table probabilities of dying, q_[x]_ (probability of dying between month *x* and month *x*+*1*), assuming a uniform distribution of deaths across age.

Second, mortality rates by age were adjusted using the most recent estimates of neonatal, infant, and under-5 mortality rates from the Inter-agency Group for Child Mortality Estimation (IGME)^6,37^ to minimize the risk of measurement errors in direct estimates of under-5 deaths based on FBH [e.g., survivor and truncation bias]^36,38^. After the previous adjustment, mortality profiles were smoothed over ages and years by using a two-dimensional P-Spline smoothing and generalized linear model, assuming that the number of deaths at a given rate are Poisson-distributed^39^.

Third, a variant of the Lee-Carter (LC)^40^ model (LLT), suitable for mortality profiles using datasets that contain multi-year gaps, was used to fit the age mortality profiles after smoothing for each country. The resulting fits were first used to generate smoothed point estimates of age-specific death rates within the 1990-2017 period.

More details about the construction of age profiles, adjustment, fit, and goodness of fit are available elsewhere^17^.

### Early mortality compression, convergence in distribution, and age-inequality

#### 1) Generalized concentration index and the potential limits of under-5 mortality decline

To measure the degree of mortality compression of U5M age profiles and age-inequality we use the Lorenz or concentration curve, denoted here as L_[0]_, and the Gini coefficient. We build on Hanada (1983)^41^ and Shkolnikov et al. (2003)^11^ who introduced the Gini coefficient as a measure of inequality of life table data. In general, the concentration curve plots the cumulative proportion of one variable (e.g., health) against the cumulative proportion of the population ranked by another variable (e.g., income). If the ranking of units of analysis (by health) is the same as the ranking of the population, the concentration and the Lorenz curve are the same. The concentration (Lorenz) curve lies below the line of 45° when poor health is concentrated in the lowest ranked group^42^. The Concentration (Gini) index (C) is then calculated as twice the area between the concentration (Lorenz) curve and the diagonal, and we define it to measure concentration of under-5 mortality as^42^:

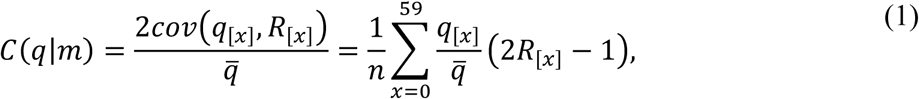

where *q*_[*a*]_ is defined as before as the probability of dying between month *x* and month *x*+*1, R*_[*x*]_ is the relative rank of the probability of dying at month *x*, 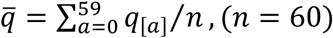, (*n* = 60) is the mean level of mortality rate in the age range of 0 to 59 months.

The Gini index as defined in equation (1) is a measure of relative inequality, and it is invariant to proportionate changes in mortality levels. This represents a limitation in the context of child mortality, particularly at advance stages of the transition where levels or changes in the neonatal mortality occur disproportionately in relation to those in other age groups (see Extended Data Fig. 3). To address this limitation, we used a generalized version of the Gini index (G_[0]_) that is obtained by multiplying C(q|m) by the mean level of mortality rate 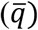. With this adjustment, G_[0]_ becomes invariant to equal improvements in mortality across age groups. The generalized Gini can then be expressed as:

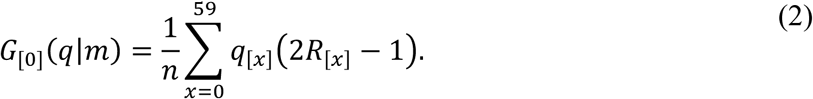

In the context of U5M, we interpret declining values of G_[0]_ as evidence of reduced inequality, convergence or early compression of under-5 deaths, with the relative mortality concentrated at ages close to 0, but declining with decreasing the mean probability of dying 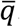.

An important feature of this index is that it is upper bounded by 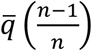,with n=60, the number of age groups. This upper limit is particularly relevant for this research because under-5 mortality is more concentrated at early ages, the Lorenz curve is below the diagonal, and although the Gini approaches 1 [more precisely 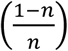] as mortality concentrates around age 0, G_[0]_ decreases with the decline of the mean probability of dying. That is, the generalized Gini index approaches the mean probability of death as the early rectangularization of mortality proceeds. In the hypothetical event of perfect rectangularization, where all deaths are concentrated at age 0 (*q*_[*x*]_ → 0, for x > 0), then the limit of G_[0]_ is proportional to the neonatal mortality level 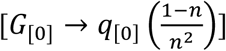, and U5M approaches *q*_[0]_ in that limit. In Figs. 1b and 3b we illustrate this hypothetical scenario—of all under-5 deaths occurring at the neonatal level—with diagonal dashed lines, that illustrate the extent of mortality convergence across countries as their neonatal and U5M rates decline at different rates. Analysis of mortality concentration was conducted using the *Stata* command ‘conindex’^43^.

#### 2) Other measures of lifespan inequality

To measure changes in age patterns of under-5 mortality and convergence in distribution towards age 0, we considered other measures of inequality that have been used in previous studies to analyze mortality of adult profiles. We first computed the minimum interval where a percentage *p* of the total under-5 deaths takes place, that we denoted *A*_*P*_. This indicator is equivalent to the C-family of indicators proposed by Kannisto^10^ for assessing compression of mortality, except that the author used it for adult mortality and our indicator always includes the age of 0 (modal age for under-5 deaths)—the age where the higher percentage of under-5 deaths occurs. For instance, while A_75_ represents the interval where 75% of under-5 deaths take place, C_75_ is the age interval where 75% of deaths in the whole population takes place (Extended Data Fig. 4).

Another indicator that we considered to account for differences across age groups and compression of U5M is the ‘conditional’ mean-age-at-death based on survival to age ‘x’ months (that we denote as [x] along this study) but death by age 5 years; we denote these by E_[x]_. Thus, E_[0]_, E_[1]_, and E_[12]_ respectively represent the ‘conditional’ mean age-at-death after surviving birth, survival to age 1 month, and survival to age 12 months. Finally, we assessed the between-individual inequality of age-at-death using the ‘conditional’ standard deviation S_[x]_ that assumes survival to age ‘x’ and death by age 5 years. We selected this indicator on the basis of theoretical^11^ and empirical^12^ findings on measures of lifespan inequality. Formally, we estimated E_[x]_ and S_[x]_ using the following expressions:

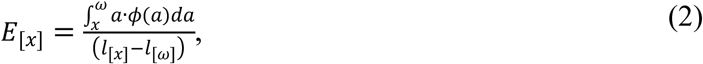

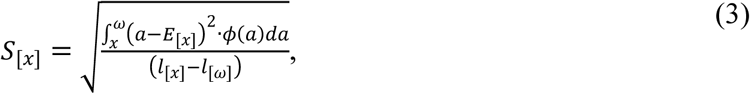

where *ω* is the oldest age in the conditional life table (59 months), *l*_[*x*]_ is the survivorship or probability of living to at least age x, and *ϕ*(*a*) = *μ*(*a*)*l*(*a*) is a density representing the probability that an individual dies at age *a* [*μ*(*a*) is the age-specific mortality rate].

As with the Gini coefficient, these three indicators (A_75,_ E[x], S[x]) represent relative measures of mean and variation and did not account for differences in the level of mortality across age groups. This was particularly problematic for analyzing age profiles of developed countries, where these indicators provided evidence of relative mortality differences that did not account for changes in absolute mortality levels across these countries, particularly at the neonatal level.

### Conditional marginal effects of income on under-5 mortality

Using information of GDP per capita—expressed in international Purchasing Power Parity (PPP) 2011 United States Dollars (USD)—and under-5 mortality for 217 countries obtained from the World Development Indicators^44^, we conducted an ecological association of mortality an national income using a scatter plot association with values from 2017. We superimposed the association from 1990 and 2017 using average values of U5M and income of countries stratified by 4 economic regions: low-income (LIC), low-middle-income (LMC), upper-middle-income (UMC), and high-income (HIC) (Fig. 3a). We further adjusted a curve that represents the prediction of under-5 mortality from an estimated fractional polynomial of income using the command ‘twoway fpfit’ from Stata. We replicated the ecological analysis using values from 1990 and with neonatal mortality rates, highlighting in color results from representative countries from each region (i.e; Rwanda from region 1; Nigeria from region 2; Gabon from region 3; and Denmark, France, Japan, and the U.S.A from region 4: Extended Data Fig. 2).

To estimate income elasticities of under-5 mortality, we fitted linear regression models with under-5 mortality as dependent variable and real GPD per capita (gdp) as independent variable, controlling for the year of observation, using the following linear specification:

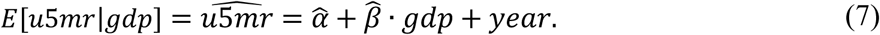

With the predicted values of under-5 mortality 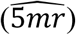 from model (7), the average elasticities are estimated as follows:

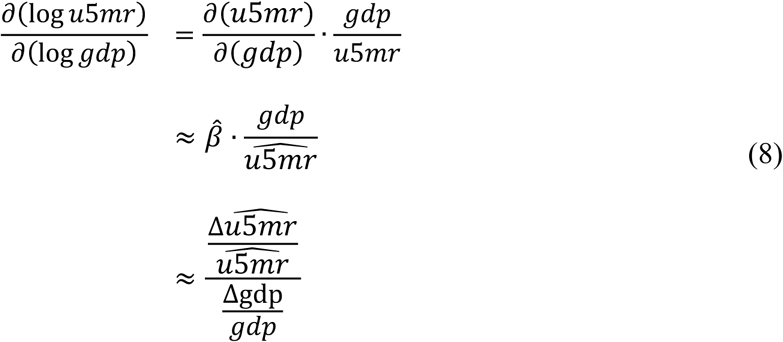

The elasticity can then be interpreted as the percentage/proportionate change in the expected mortality for a percentage/proportionate change in the GDP per capita. Elasticities were calculated in a yearly-basis relative to the median GDP per capita value of the corresponding year and region using a regression approach and the post-estimation command ‘eyex’ from *Stata*^45^.

### Strengths and limitations

Our study builds on previous research that analyzed inequality in the length of life and convergence of adult mortality^11,12,46^. Using under-5 mortality profiles from narrow-age groups from Denmark, France, Japan, and the USA from publicly available data, and from 31 SSA countries with data constructed by Mejía-Guevara et al. (2019)^17^ for the forecasting of mortality rates and the assessment of the SDG-3 by 2030, this study is—to our knowledge—the first to use this kind of mortality patterns to explore the potential limit of child survival, by assessing compression, convergence and under-5 age-inequality. In particular, the age profiles of under-5 mortality for high-income countries allowed us to investigate potential limits in child survival, as data for the 4 developed countries in our sample included the most recent trends of under-5 mortality patterns, except for Denmark where additionally relied on annual information as far as 1901, allowing a broader analysis of the mortality transition of under-5 deaths during the last 120 years. We investigated the extent of early compression, age-inequality and variation in under-5 mortality, and we found important differences in the patterns of change across countries over time, with a positive association between the rates of mortality change and age-inequality. We identified Japan as the leading country in the under-5 mortality convergence, not only due to its lowest neonatal and under-5 mortality levels, but also because the early compression and the least inequality of mortality patterns that it has achieved in the most recent years.

To highlight differences in the evolution of age-patterns, we examine the youngest ages at which 75% of under-5 mortality takes place in a country that we denote by A_75_ (Extended Data Fig. 4). In 1980, 75% of deaths in Japan occurred within the first 12 months of life and that percentage remained around that level because its sharp decline in neonatal mortality relative to other ages, in contrast with that observed in the USA over the past 50 years, where they occurred within the first 5 months because the relative importance of neonatal mortality remained the same. With much higher levels of mortality at all ages and despite being on track to meet the SDG-3 of child mortality reduction(13), Rwanda and Senegal will expect to exhibit mortality profiles comparable to those observed in Denmark and France more than 50 years ago (Fig. 2 and Extended Data Fig. 1), and with larger neonatal levels. The gap is even larger for several SSA countries, like Chad and Nigeria (Extended Data Table 1).

We identified salient differences across SSA countries in terms of distribution of deaths by age and time, convergence, and inequality in mortality reduction by age; particularly after stratifying countries on the basis of their achieved ARR between 1990 and 2016. On the one hand, countries with an ARR below 3.2% experienced a slower convergence process both in terms of mean and distribution of age-at-death. Mortality reductions occurred at different rates across under-5 ages, the mean and dispersion of age-at-death remained constant or have increased in most countries, which is in line with results from previous studies^47^, showing unusual high values of _4_q_1_ (the probability of children dying between age 1y and 5y) relative to _1_q_0_ (the probability of dying during the first year of life) in several countries from SSA, that may be attributable to true epidemiological patterns (e.g., disease environments characterized by high prevalence of infectious diseases—malaria, measles, and diarrhea) rather than to data quality issues as similar patterns have been found in previous studies based on Demographic Surveillance Systems (DSS) data^47,48^. On the other hand, our findings revealed substantive evidence of mortality convergence, both in terms of mean and distribution, for countries that have reached higher ARR (≥3.2%), including countries that have met the MDG-4—e.g., Ethiopia, Malawi, Niger, Tanzania, and Rwanda^49^. However, when compared with the age profiles of highly developed countries, even those like Rwanda and Senegal which are on track to achieve the SDGs by 2030^17^, face a difficult path ahead to achieve further reductions of child mortality according to our estimates. The relationship between national income and under-5 mortality is in line with previous evidence highlighting income as a key predictor of mortality gaps across economic regions^18^. We included that evidence as our analysis is based on mortality distributions from two groups of countries situated at both extremes of the income distribution and stages of the mortality transition. The analysis reveal that despite that association, it is unclear whether increases in income are enough to overcome the mortality gaps between countries in advanced transitions, for example between Japan and the USA, or between countries in the earliest transition, even though the important reductions achieved in countries like Rwanda during the past 25 years may not be attributed to income growth changes but to the implementation of key policy interventions^50^. It is also unclear, and our data are not suitable to response whether further reductions in mortality in poor settings may be achievable without substantial improvements in economic development, or the extent to which the USA would be able to reach similar levels as Japan despite its more advanced economic position.

The scope of this study is limited for several reasons. Although our data for Denmark, France, Japan, and the USA were obtained from high quality vital registration records that represent diverse world regions, we only had information from those 4 developed countries. For SSA, we relied on full-life histories from survey data, which are subject to several sources of error; as vital statistics for this group of countries are inexistent or deficient in the countries included in our study^51^. Second, we used different measures of inequality to improve the robustness of our analysis as they differ in their underlying properties and sensitivity to age-specific mortality change^14^. Third, because of the nature of the survey data, we cannot infer causality neither in the association of the average decline of mortality and age-inequality nor between mortality and national income, and further research should explore the underlying risk factors, social determinants, and true epidemiological patterns behind under-5 mortality change by age, particularly in the SSA region.

## Extended Data

**Extended Data Fig. 1.**
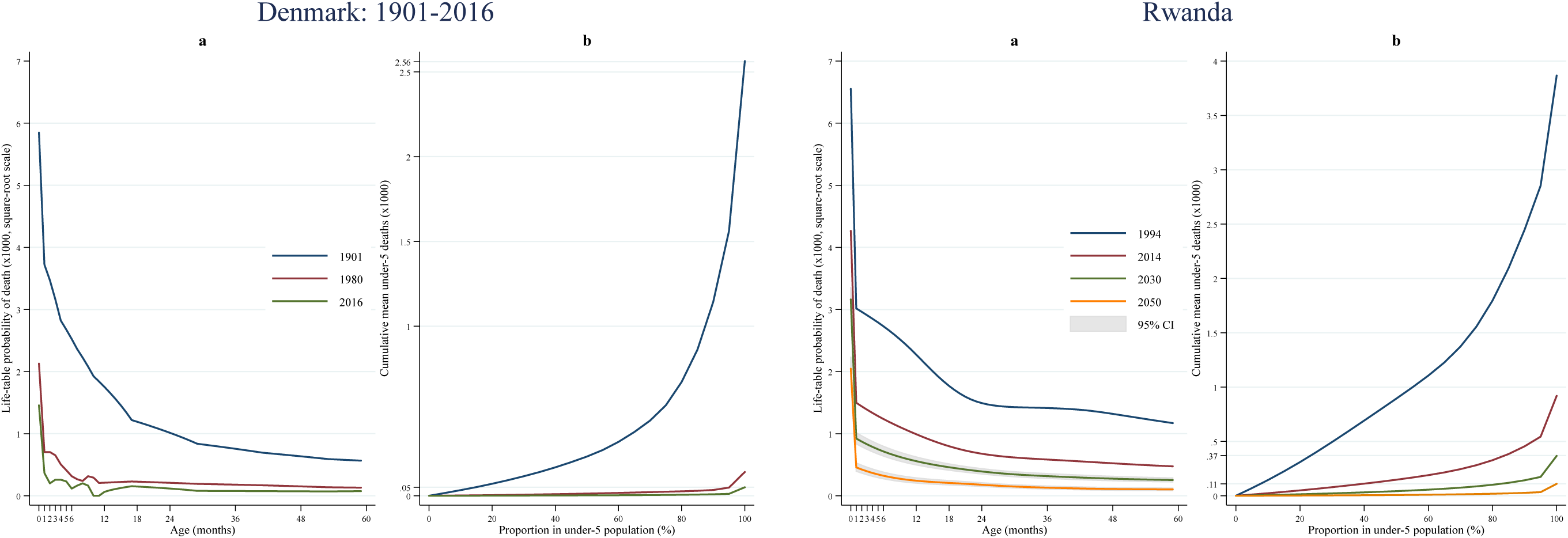
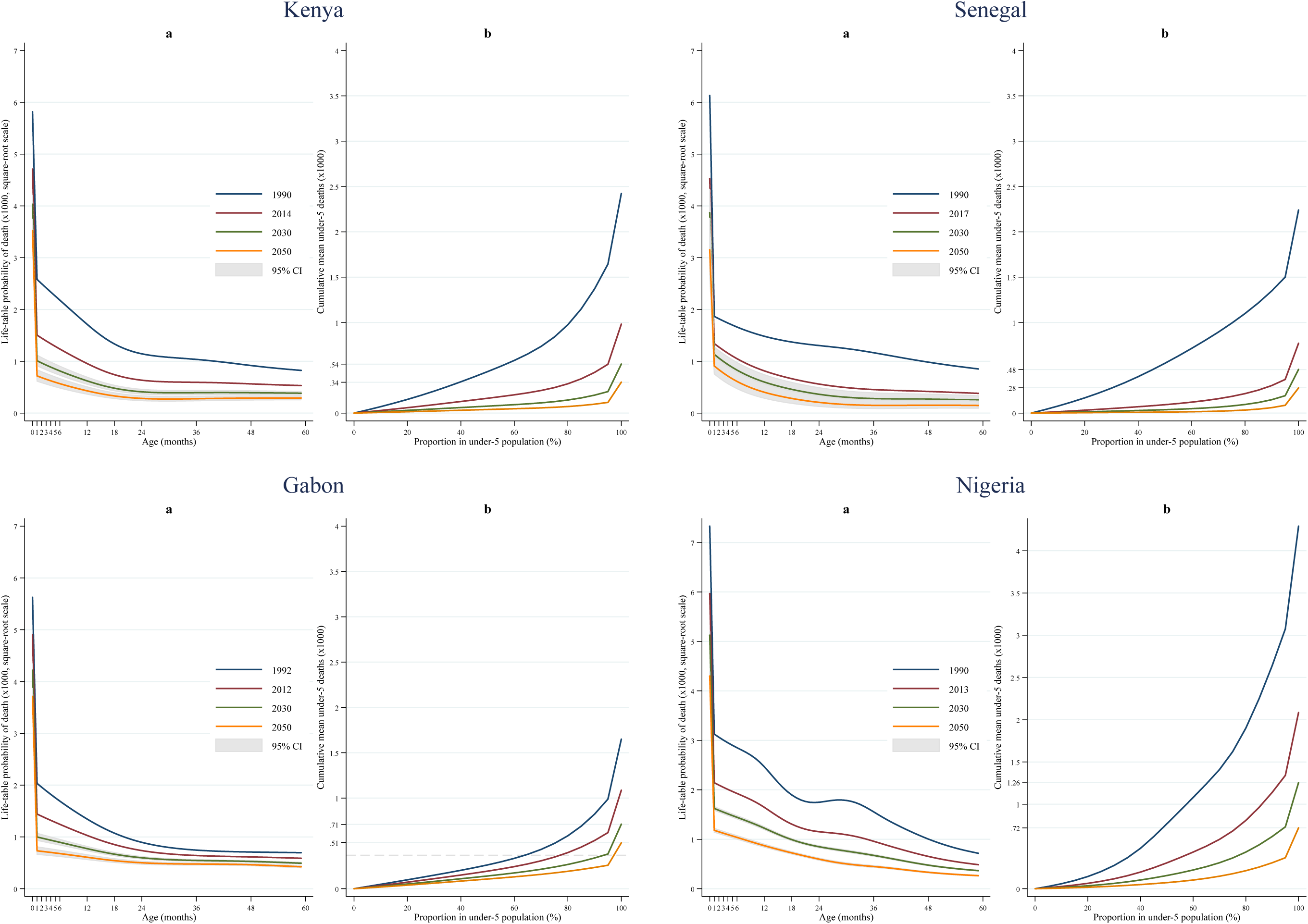
Age patterns of under-5 deaths and inequality in the age at death. **a**, Detailed age-at-death distribution of under-5 deaths in Denmark (1901-2016) and four selected countries from SSA (Gabon, Kenya, Nigeria, and Senegal). Age is measured in months in the x-axis and life-table probabilities of dying are represented in the y-axis (x1000 in square-root scale). Forecasts of mortality age profiles were conducted for countries in SSA by 2030 and 2050. **b**, Generalized Lorenz curves in the same selected countries showed in **a)**. Generalized Lorenz curves are constructed as the product of the standard Lorenz curve (Extended Data Fig. 1) and the mean mortality rates across ages (measured in months). It therefore illustrates the cumulative percentage of under-5 population (x-axis), ranked by the level of mortality at each age, and graphed against the cumulative mean under-5 deaths (y-axis) (Supplementary Methods and Extended Data Table1).

**Extended Data Fig. 2.**
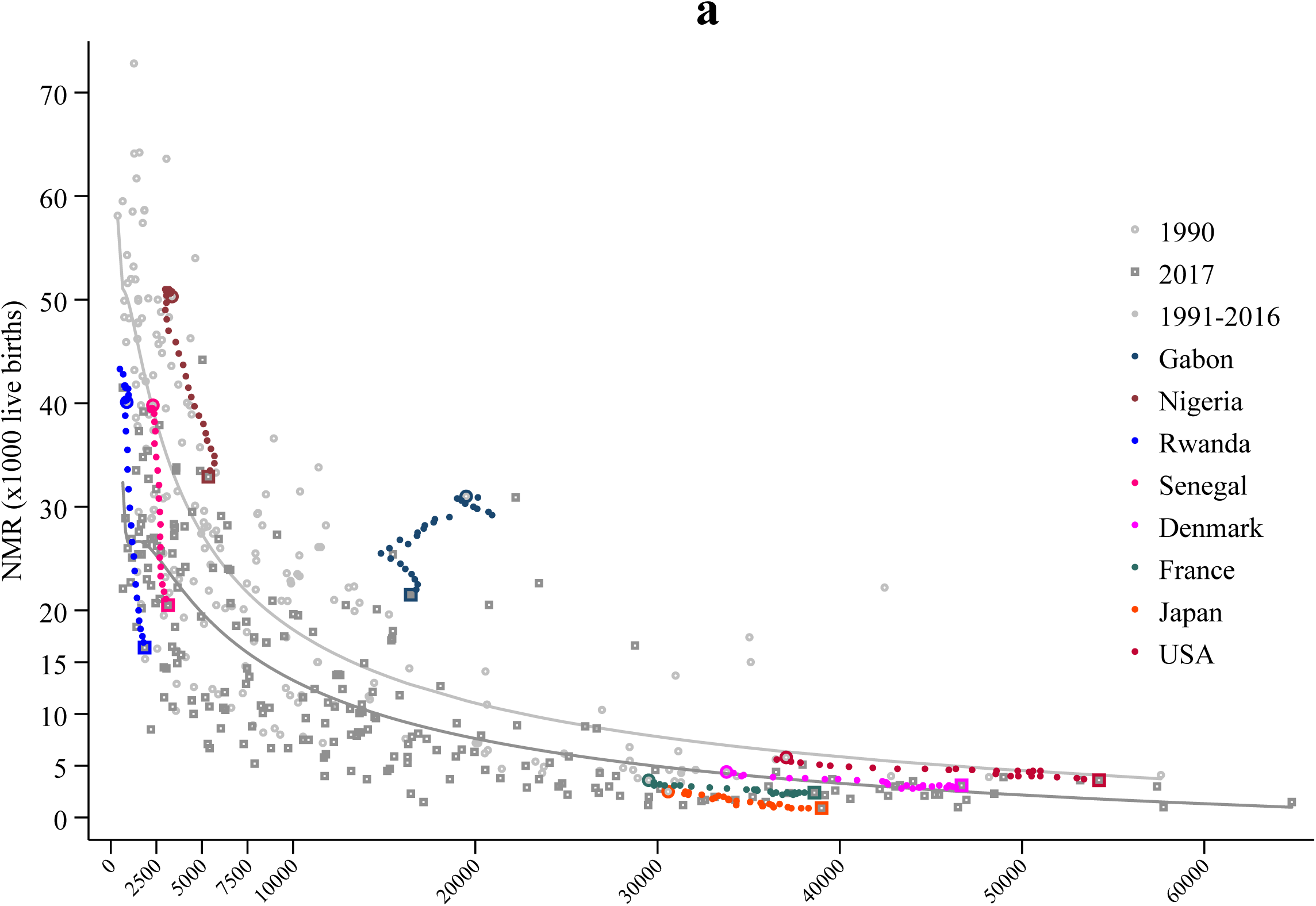
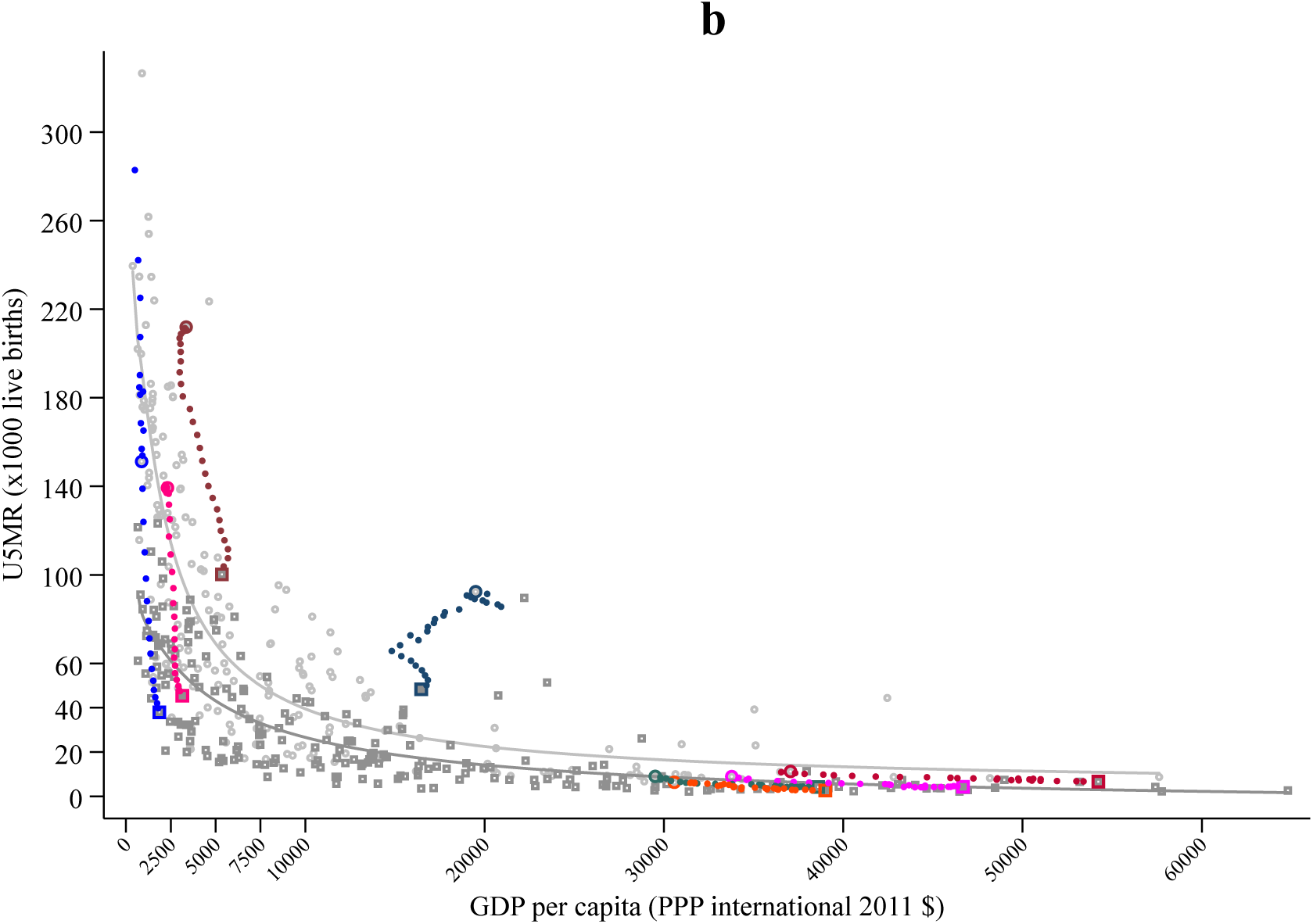
Worldwide association of neonatal and under-5 mortality and economic development. **a**, Country-wise association of GDP per capita income and neonatal mortality rates in 1990 and 2017. **b**, Country-wise association of GDP per capita income and under-5 mortality rates in 1990 and 2017. GDP per capita is expressed in international 2001 PPP USD, mortality rates as deaths per 1,000 live births, and yearly average values are shown for the period between 1990 and 2017 by economic region (in color). The adjusted curves represent the prediction of neonatal or under-5 mortality from an estimated fractional polynomial of income. Data are from World Development Indicators, retrieved online from The World Bank (2019), which included 217 countries worldwide (8 countries with GDP above 70,000 are not displayed in panel **b**), classified in 4 economic regions: low-income (LIC—pink), low-middle-income (LMC—maroon), upper-middle-income (UMC—blue), and high-income (HIC—orange) (Supplementary Methods).

**Extended Data Fig. 3.**
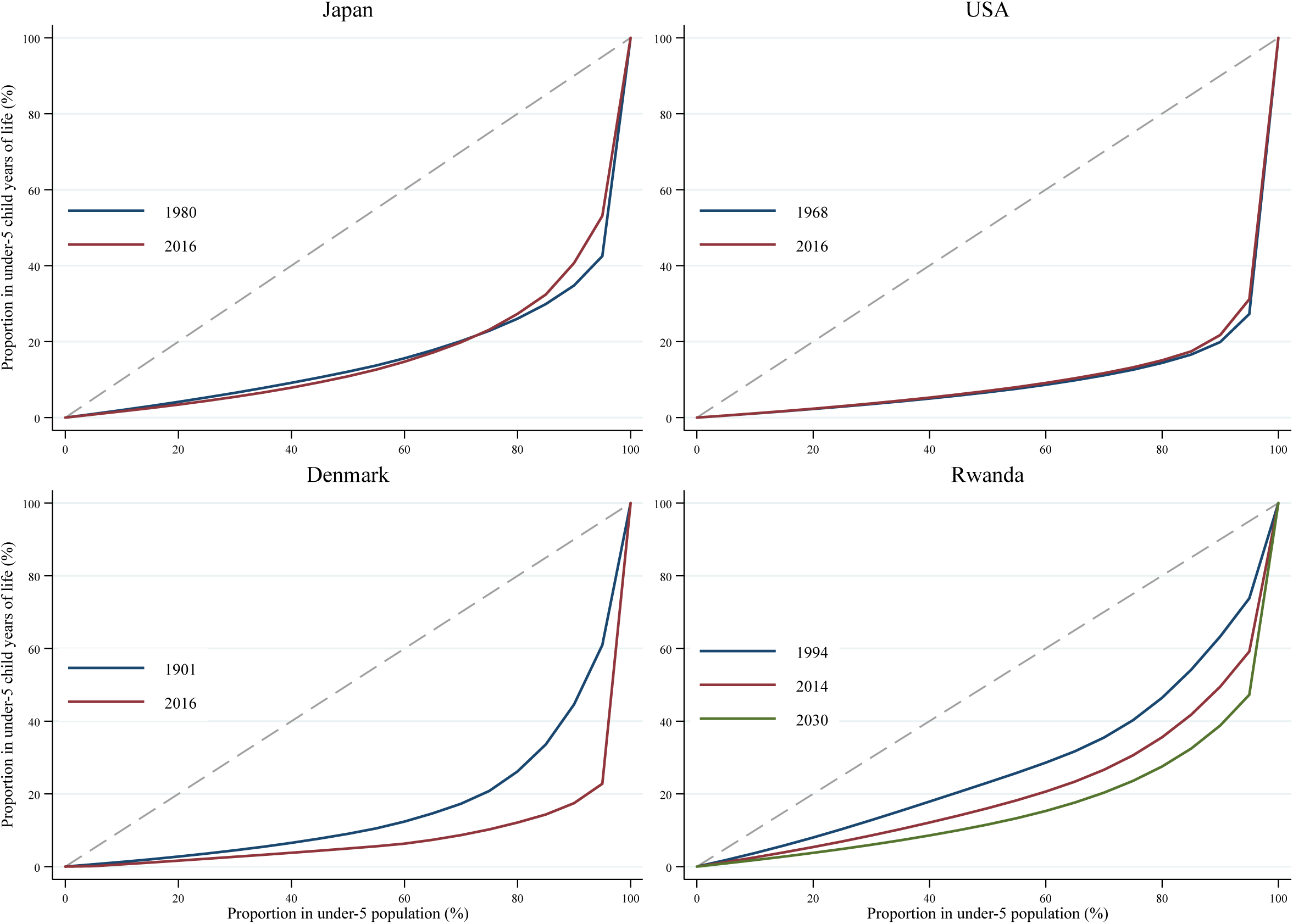
Under-5 inequality in the age at death. Standard Lorenz curves for the first and the latest observed years in Denmark (1901, 2016), Japan (1980, 2016), the U.S.A. (1968, 2016), and Rwanda (1994, 2014, 2030). The Lorenz curve measures age inequality in the age at death distribution of under-5 deaths, and indicate less inequality if they move away (to the right) from the 45° line that in the context of child mortality represents mortality evenly distributed across ages. Two times the area gap between the 45° line and the Lorenz curve is known as the Gini coefficient (G_[0]_), which increases as under-5 inequality declines (until certain level where the relative importance of neonatal mortality declines relative to that at other ages, as occurring in the case of Japan or the USA) (Supplementary Methods and Extended Data Table1).

**Extended Data Fig. 4.**
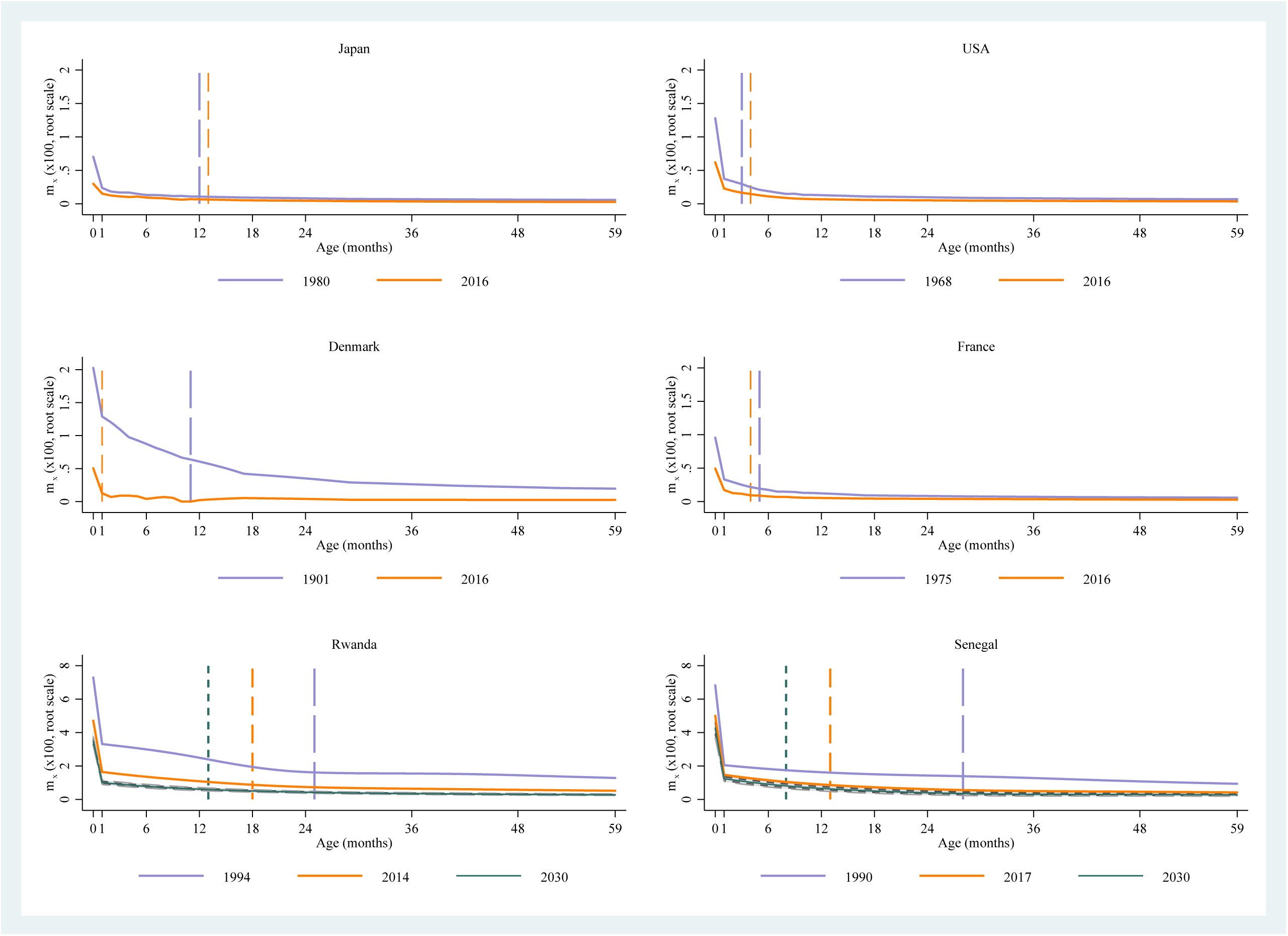
Age distribution and compression of under-5 mortality. Change in mortality rates by age and the upper quartile of the conditional distribution of age at death (A_75_) between the first and latest observed year, and projected by 2030 (the target year for the SDG-3 of child mortality reduction) in selected countries: Denmark, France, Japan, USA, Rwanda, and Senegal. These countries were selected to represent different trajectories in developed countries and to illustrate trajectories of the most successful countries in SSA in terms of under-5 mortality reduction within the past 25 years, like Rwanda and Senegal advancing rapidly and likely to reach the SDD-3 by 2030, but with relatively high neonatal mortality as compared to Denmark and France, countries in a very advanced mortality transition and low child mortality. Vertical lines represent the age where 75% of under-5 deaths took place in the respective year. For instance, in Rwanda, 75% of under-5 deaths occurred during the first 2 years of life in 1994, before 18 months in 2014, and they would occur before 12 months in 2030; whereas in Denmark, the 12 months of life concentrated the same 75% of under-5 deaths around 1901 (130 years ago). Data are from 31 DHS surveys and publicly available mortality records from Denmark, France, Japan, and the USA (Supplementary Methods).

**Extended Data Table 1.** Income, mortality, and age-at-death indicators of compression and inequality.

| Country | Year | | T | GDP (2011 PPP\$) | | | Neonatal Mortality | | | Under-5 Mortality | | | A<sub>75</sub> | | E<sub>[0]</sub> | | S<sub>[0]</sub> | | G<sub>[0]</sub> | |
| --- | --- | --- | --- | --- | --- | --- | --- | --- | --- | --- | --- | --- | --- | --- | --- | --- | --- | --- | --- | --- |
|  | (1) | (2) |  | (1) | (2) | AGR | (1) | (2) | ARR | (1) | (2) | ARR | (1) | (2) | (1) | (2) | (1) | (2) | (1) | (2) |
| **Low-income** | 1990 | 2017 |  |  |  |  |  |  |  |  |  |  |  |  |  |  |  |  |  |  |
| Benin | 1990 | 2012 | 24 | 1463 | 1856 | 1.1 | 45 | 35 | 1.1 | 177 | 108 | 2.2 | 24 | 21 | 14.3 | 13.2 | 16.4 | 16.3 | 1.5 | 1.0 |
| Burkina Faso | 1990 | 2010 | 22 | 844 | 1423 | 2.6 | 44 | 31 | 1.8 | 199 | 116 | 2.7 | 27 | 24 | 15.9 | 14.6 | 16.3 | 16.0 | 1.5 | 1.0 |
| Burundi | 1990 | 2017 | 11 | 1032 | 668 | -1.6 | 40 | 22 | 2.2 | 179 | 62 | 3.9 | 23 | 16 | 14.7 | 10.6 | 16.7 | 14.6 | 1.5 | 0.7 |
| Chad | 1990 | 2015 | 22 | 1110 | 2067 | 2.5 | 50 | 36 | 1.3 | 215 | 131 | 2.0 | 26 | 25 | 15.7 | 14.5 | 16.5 | 16.7 | 1.7 | 1.1 |
| DR of Congo | 1991 | 2014 | 22 | 1224 | 785 | -1.9 | 42 | 30 | 1.4 | 193 | 102 | 2.8 | 20 | 13 | 13.1 | 10.1 | 15.1 | 13.9 | 1.8 | 1.1 |
| Ethiopia | 1990 | 2016 | 28 | 652 | 1608 | 3.5 | 60 | 30 | 2.7 | 204 | 64 | 4.4 | 26 | 16 | 14.4 | 10.7 | 17.4 | 15.8 | 1.8 | 0.7 |
| Gambia | 1993 | 2013 | 6 | 1504 | 1585 | 0.3 | 48 | 30 | 2.3 | 154 | 73 | 3.7 | 33 | 22 | 17.8 | 12.7 | 18.8 | 16.6 | 1.0 | 0.7 |
| Guinea | 1990 | 2012 | 24 | 1412 | 1684 | 0.8 | 67 | 28 | 3.9 | 240 | 103 | 3.8 | 22 | 20 | 13.4 | 12.9 | 15.8 | 15.0 | 2.2 | 1.0 |
| Liberia* | 1991 | 2011 | 12 | 1292 | 1139 | -1.2 | 56 | 29 | 3.2 | 257 | 96 | 4.8 | 16 | 15 | 11.6 | 10.4 | 14.2 | 13.5 | 2.7 | 1.0 |
| Malawi | 1990 | 2016 | 23 | 744 | 1084 | 1.5 | 52 | 23 | 3.1 | 256 | 71 | 4.8 | 21 | 21 | 13.8 | 12.7 | 15.1 | 15.7 | 2.4 | 0.7 |
| Mali | 1990 | 2012 | 24 | 1272 | 1805 | 1.6 | 67 | 39 | 2.4 | 252 | 126 | 3.1 | 27 | 27 | 15.6 | 15.1 | 16.4 | 17.3 | 2.0 | 1.0 |
| Mozambique | 1990 | 2011 | 23 | 379 | 955 | 4.5 | 60 | 31 | 3.1 | 261 | 100 | 4.5 | 18 | 16 | 12.2 | 11.4 | 15.1 | 15.1 | 2.7 | 1.0 |
| Niger | 1990 | 2012 | 24 | 895 | 863 | -0.2 | 55 | 30 | 2.8 | 345 | 114 | 4.9 | 30 | 27 | 18.0 | 15.5 | 15.7 | 16.3 | 2.4 | 0.9 |
| Rwanda | 1994 | 2014 | 22 | 497 | 1633 | 6.1 | 43 | 18 | 4.2 | 208 | 54 | 6.5 | 25 | 18 | 15.3 | 12.0 | 16.6 | 15.6 | 1.7 | 0.5 |
| Senegal | 1990 | 2017 | 28 | 2314 | 3143 | 1.1 | 38 | 21 | 2.2 | 126 | 45 | 3.7 | 28 | 13 | 15.9 | 9.4 | 17.1 | 14.5 | 1.0 | 0.5 |
| Sierra Leone | 1993 | 2013 | 6 | 1052 | 1655 | 2.3 | 52 | 38 | 1.6 | 252 | 138 | 3.0 | 21 | 14 | 13.5 | 9.7 | 15.2 | 11.7 | 2.4 | 1.6 |
| Togo | 1990 | 2014 | 13 | 1294 | 1406 | 0.3 | 43 | 27 | 1.9 | 150 | 80 | 2.6 | 22 | 20 | 13.6 | 12.6 | 16.1 | 16.1 | 1.4 | 0.8 |
| Uganda | 1990 | 2016 | 24 | 769 | 1689 | 3.1 | 35 | 21 | 1.9 | 186 | 53 | 4.7 | 21 | 14 | 14.1 | 9.7 | 15.4 | 13.7 | 1.6 | 0.6 |
| Tanzania | 1990 | 2016 | 19 | 1473 | 2584 | 2.2 | 37 | 22 | 2.1 | 170 | 57 | 4.2 | 21 | 16 | 13.9 | 10.5 | 15.7 | 14.5 | 1.5 | 0.6 |
| Zimbabwe | 1990 | 2015 | 16 | 2889 | 2197 | -1.1 | 29 | 24 | 0.8 | 103 | 75 | 1.2 | 24 | 21 | 14.1 | 13.0 | 16.5 | 16.2 | 0.9 | 0.7 |
| **Lower-middle-income** |  |  |  |  |  |  |  |  |  |  |  |  |  |  |  |  |  |  |  |  |
| Angola | 1991 | 2016 | 7 | 4538 | 6265 | 1.3 | 54 | 30 | 2.4 | 232 | 85 | 4.0 | 25 | 20 | 14.7 | 12.4 | 16.7 | 16.3 | 2.0 | 0.8 |
| Cameroon | 1990 | 2011 | 23 | 3020 | 2969 | -0.1 | 34 | 29 | 0.9 | 143 | 106 | 1.4 | 21 | 20 | 13.5 | 12.9 | 15.4 | 15.1 | 1.3 | 1.0 |
| Congo | 1993 | 2012 | 13 | 4928 | 5271 | 0.4 | 34 | 22 | 2.4 | 126 | 71 | 2.9 | 20 | 19 | 13.1 | 12.0 | 15.8 | 15.0 | 1.2 | 0.7 |
| Cote d'Ivoire | 1990 | 2012 | 15 | 3194 | 2712 | -0.7 | 52 | 37 | 1.6 | 164 | 112 | 1.7 | 16 | 16 | 11.2 | 11.2 | 14.8 | 14.9 | 1.7 | 1.2 |
| Ghana | 1991 | 2014 | 13 | 1966 | 3828 | 2.9 | 40 | 27 | 1.7 | 117 | 59 | 2.9 | 21 | 15 | 12.9 | 9.7 | 16.1 | 14.2 | 1.1 | 0.7 |
| Kenya | 1990 | 2014 | 15 | 2380 | 2753 | 0.6 | 34 | 22 | 1.7 | 136 | 57 | 3.5 | 21 | 17 | 13.4 | 11.6 | 15.9 | 16.1 | 1.2 | 0.6 |
| Lesotho | 1995 | 2010 | 11 | 1628 | 2366 | 2.5 | 42 | 42 | 0.1 | 111 | 116 | -0.3 | 15 | 15 | 10.4 | 10.6 | 15.0 | 15.0 | 1.2 | 1.2 |
| Nigeria | 1990 | 2013 | 25 | 3359 | 5492 | 2.2 | 54 | 36 | 1.8 | 229 | 118 | 2.8 | 21 | 19 | 13.1 | 11.8 | 14.3 | 14.1 | 2.2 | 1.2 |
| **Upper-middle-income** |  |  |  |  |  |  |  |  |  |  |  |  |  |  |  |  |  |  |  |  |
| Zambia | 1990 | 2014 | 23 | 2342 | 3633 | 1.8 | 37 | 24 | 1.9 | 180 | 68 | 4.0 | 22 | 17 | 13.8 | 11.2 | 14.9 | 14.9 | 1.6 | 0.7 |
| Gabon | 1992 | 2012 | 10 | 19006 | 16159 | -0.8 | 32 | 24 | 1.4 | 95 | 63 | 2.0 | 18 | 20 | 11.9 | 12.5 | 15.7 | 16.5 | 0.9 | 0.6 |
| Namibia | 1992 | 2007 | 7 | 6270 | 8098 | 1.7 | 27 | 20 | 2.0 | 76 | 69 | 0.6 | 17 | 18 | 11.2 | 12.2 | 15.0 | 15.3 | 0.8 | 0.7 |
| **High-income\*** |  |  |  |  |  |  |  |  |  |  |  |  |  |  |  |  |  |  |  |  |
| Denmark | 1901 | 2016 | 116 | 33786 | 45991 | 1.2 | 34 | 2 | 2.4 | 143 | 3 | 3.3 | 11 | 1 | 8.8 | 4.7 | 12.3 | 11.2 | 1.7 | 0.0 |
| France | 1975 | 2016 | 42 | 29515 | 38063 | 1.0 | 8 | 2 | 3.2 | 14 | 3 | 3.3 | 5 | 4 | 6.3 | 5.9 | 12.4 | 12.3 | 0.2 | 0.0 |
| Japan | 1980 | 2016 | 37 | 30582 | 38283 | 0.9 | 4 | 1 | 4.6 | 8 | 2 | 3.6 | 12 | 13 | 9.0 | 9.5 | 14.7 | 13.8 | 0.1 | 0.0 |
| USA | 1968 | 2016 | 49 | 37062 | 53399 | 1.4 | 14 | 3 | 3.0 | 21 | 6 | 2.7 | 3 | 4 | 5.2 | 5.6 | 11.7 | 11.9 | 0.3 | 0.1 |

\* GDP for Liberia corresponds to 2001 and to 1990 for Denmark, France, Japan, and the U.S.A.
Complete list of countries classified by income group, and indicators of income and mortality for the initial and end year of analysis: GDP per capita levels (in international PPP 2011\$) (GDP for Liberia corresponds to 2001 and to 1990 in Denmark, France, Japan, and the USA), and average growth rate (AGR); neonatal and under-5 mortality rates, and average rates of reduction (ARR); upper quartile of the conditional distribution of age at death ( $A_{75}$ ); mean ( $E_{[0]}$ ) and standard deviation ( $S_{[0]}$ ) of age at death; and Generalized Gini Index ( $G_{[0]}$ ).

## Supplementary Information

**Fig. S1.**
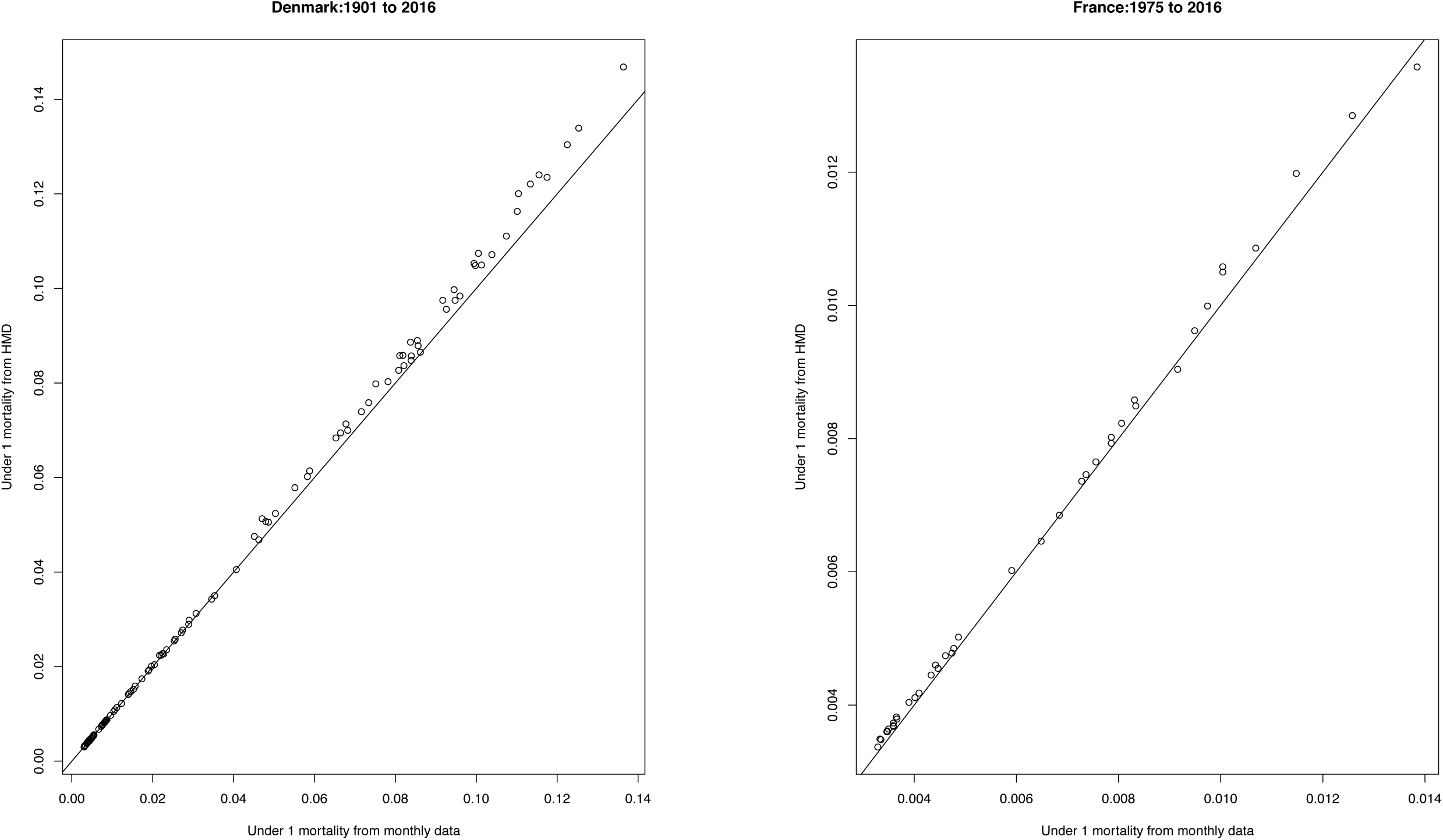
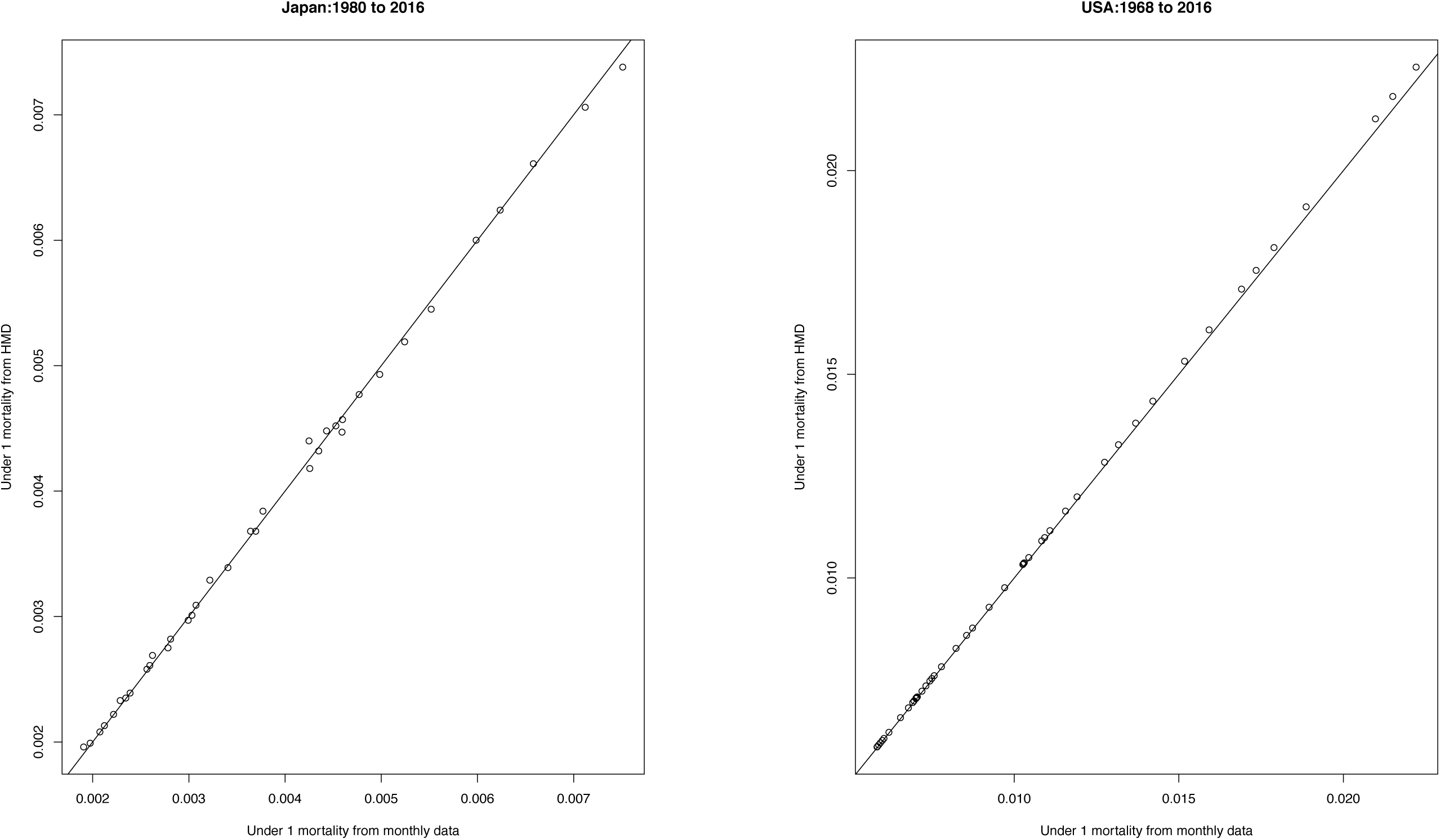
Assessment of infant mortality fit from monthly-based age profiles and infant mortality from the Human Mortality Database (HMD) from Denmark, France, Japan, and the USA (Supplementary Methods)

